# Snapshots of Internal Protein Crystal Architecture at the Nanoscale

**DOI:** 10.64898/2026.01.16.700027

**Authors:** Shervin S. Nia, Ambarneil Saha, Jung Cho, Z. Hong Zhou, Peter Ercius, Matthew Mecklenburg

## Abstract

Macromolecular crystallography has historically inferred models of internal crystal architecture from reciprocal-space measurements of Bragg reflections. Nevertheless, direct real-space visualization of crystallographic disorder remains elusive, particularly at the nanoscale. Using a 15-nanometer probe, here we apply both ambient-temperature and cryogenic four-dimensional scanning transmission electron microscopy (4D–STEM) to map the topography of coherently diffracting domains (CDDs) in lysozyme and myoglobin microcrystals at length scales 100× finer than conventional X-ray and electron beams. Virtual dark-field reconstructions from individual Bragg reflections reveal that each peak arises from spatially distinct subvolumes. Subdomain analysis demonstrates that these CDDs behave as orientationally monolithic units analogous to single mosaic blocks. Under sustained irradiation, protein CDDs undergo continuous rearrangements spanning several micrometers of internal movement. Furthermore, pinpoint high-dose “impact crater” experiments reveal delocalized radiolytic damage propagating hundreds of nanometers from primary irradiation sites, behavior similar to small-molecule crystals but amplified in both rate and magnitude. Together, these results establish macromolecular microcrystals as dynamic assemblies whose internal architecture continuously reorganizes during irradiation, laying the foundation for realistic models of mosaicity directly informed by both real-space and reciprocal-space observations.

## Introduction

X-ray crystallography has uniquely defined the inception and trajectory of structural biology (1), delivering the first landmark structures of macromolecules such as proteins (2) and nucleic acids (3). Successful macromolecular structure determination depends critically on crystal quality, particularly on the degree of long-range order that governs diffraction to higher Bragg angles. Nevertheless, macromolecular crystals typically contain significant imperfections that limit their diffracting power, often thwarting efforts aimed at high-resolution structure elucidation (4).

Mapping the exact spatial distribution of these imperfections has historically proven challenging. Micron-scale X-ray beams (5) average signal over vast numbers of unit cells, obscuring nanoscale heterogeneities. This limitation has spurred X-ray crystallographers to work backwards, inferring the presence of defects in real-space crystal architecture by analyzing reciprocal-space measurements of Bragg reflections (4). Specifically, the prevailing model of mosaicity (6; 7) conceptualizes crystals as patchworks of misaligned “mosaic blocks” whose angular misorientation in real space produces peak broadening effects (8) in reciprocal space (Fig. 1A). However, a typical X-ray diffraction experiment cannot directly image the size, shape, and distribution of these putative mosaic blocks, and mosaicity makes no attempt to model the boundaries or discontinuities separating miniature crystallites (9). In other words, a real-space perspective is lacking. Consequently, reciprocal-space signatures such as distorted Bragg peak profiles have provided the only major window into crystal imperfections, leaving our understanding of the underlying organization of nanoscale coherently diffracting domains (CDDs) underdeveloped.

**Figure 1.**
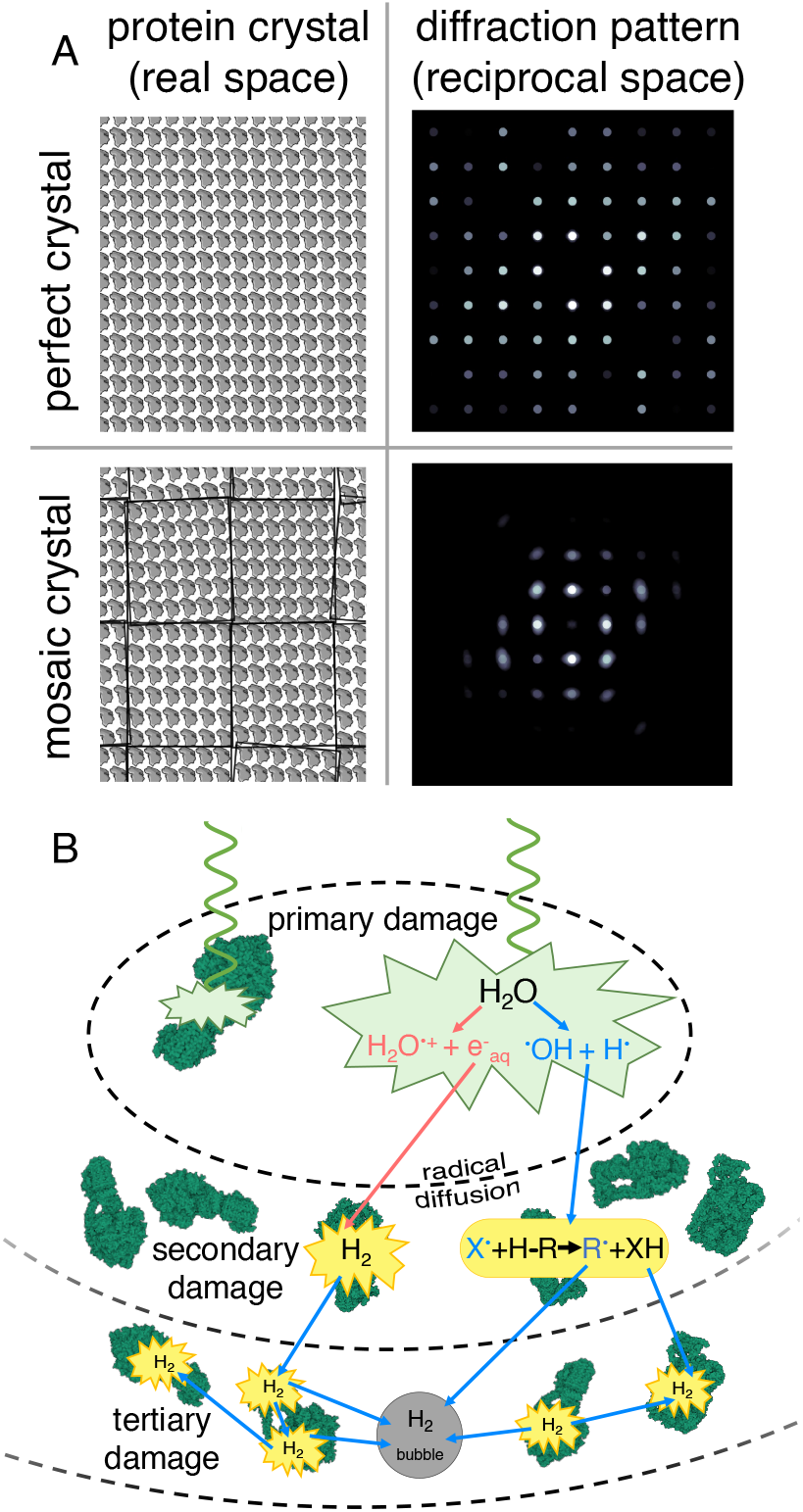
Models of mosaicity and radiation damage. (A) Schematic representations of a perfect, monolithic crystal (top row) and an imperfect, mosaic crystal (bottom row) alongside corresponding diffraction patterns (right column) simulated by taking the Fourier transform of the images on the left. The mosaic crystal contains several angularly misoriented subregions, introducing peak broadening effects in the Fourier transform. (B) Schematic representation of electron beam-induced radiolytic damage, where incident quanta (green wave) scatter inelastically within the specimen (the primary inelastic collision, represented by the green polygon), resulting in ionization, electronic excitation, secondary electron emission, and subsequent radiolytic cleavage of covalent bonds into free radicals. These reactive radiolytic fragments diffuse away from the parent bond and inflict cascades of secondary damage via downstream radical chemistry (yellow), breaking distal bonds further removed from the point of origin. Over time, many such events lead to global decay of structural integrity at all length scales (tertiary damage). A visible symptom of tertiary damage is the formation of dihydrogen gas bubbles.

The theoretical framework for understanding crystal imperfections becomes clearer when contrasting ideal and real crystals. Perfect, monolithic crystals yield identical diffraction patterns regardless of where they were sampled in real space, producing sharp, pointlike Bragg peaks (Fig. 1A). Conversely, imperfect macro-molecular crystals exhibit spatial variations in diffracting power and orientation that originate from irregularities in the underlying CDD topography. Crucially, these domains do not remain static under irradiation, as electron or X-ray beam-induced radiolytic damage continually degrades long-range order during data collection— thus broadening Bragg peaks and increasing mosaicity (10; 11). Radiolysis is initiated via core- or valence-shell ionization events that break covalent bonds, generating low-energy secondary electrons and mobile free radicals (12; 13; 14; 15). These reactive species then proceed to travel through the specimen once formed (Fig. 1B), inflicting more damage through cascades of secondary and tertiary reactions (16; 17). While many conventional measurements have documented this deterioration by monitoring the fading or broadening of diffraction spots, the corresponding real-space evolution of macromolecular crystal architecture during radiolysis is poorly understood. Furthermore, the fundamental length scale at which Bragg diffraction transitions from monolithic to mosaic—effectively, the coherence length of individual mosaic blocks—remains inaccessible to measurement by broad-beam methods.

Here we employ four-dimensional scanning transmission electron microscopy (4D–STEM; Fig. 2A) to bridge these gaps (18; 19; 20). By acquiring independent diffraction patterns at an array of probe positions separated by nanometer-scale steps, 4D–STEM generates dual-space datasets that simultaneously encode real- and reciprocal-space information at length scales matching expected mosaic block dimensions (8). We build on previous studies conducted using synchrotron X-ray topography (9; 21) and dark-field transmission electron microscopy (TEM) imaging (22), both of which have yielded valuable insights into macro-molecular crystal architecture. Our approach is differentiated from prior X-ray work by the significantly smaller probe size achievable using STEM, enabling spatial resolution 100× finer than comparable X-ray topographs. Furthermore, 4D–STEM is distinguished from prior EM work by its ability to reconstruct both dark-field images and selected-area diffraction patterns (Fig. 2B) using customizably shaped virtual apertures (23). In contrast, conventional TEM relies on manually maneuvering fixed physical apertures to chase rapidly decaying Bragg signal during the experiment itself. 4D–STEM permits direct visualization of exactly which nanoscale regions in real space produced specific Bragg reflections (Fig. 2C-F).

**Figure 2.**
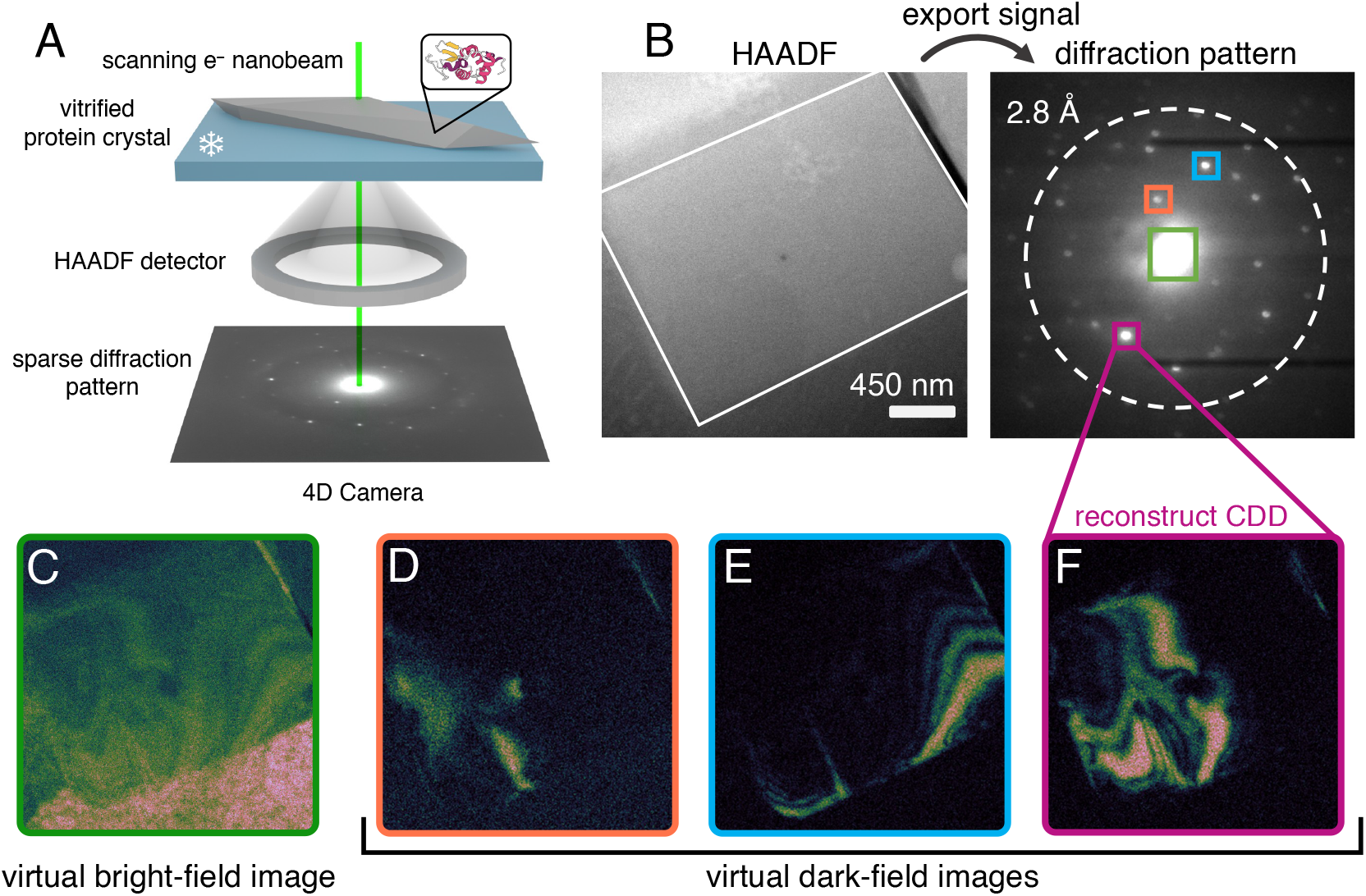
4D–STEM data collection and processing. (A) Schematic of a standard cryo-4D–STEM experiment, with a scanning electron nanobeam impinging on a vitrified protein microcrystal. A separate high-angle annular dark-field (HAADF-STEM) image is acquired on a monolithic detector, whereas coherent Bragg signal is acquired on the 4D Camera operating at 87000 frames per second. (B) Workflow for generating a selected-area diffraction pattern by masking a specific subregion (white border; virtual selected-area aperture) of a simultaneously acquired HAADF-STEM image. (C) Virtual bright-field image reconstructed by masking the unscattered beam, marked by the green square (virtual objective aperture) in (B). (D-F) Virtual dark-field images reconstructed by masking specific Bragg reflections, marked by the orange (D), blue (E), and magenta (F) squares in (B), respectively.

Using a custom-built 87 kHz frame-rate direct electron detector (24), here we apply both ambient-temperature and cryogenic 4D–STEM to interrogate the internal architecture of microcrystals formed by the model proteins lysozyme and myoglobin. We find that individual Bragg reflections arise from spatially distinct, often non-overlapping CDDs. Subdividing these CDDs into even smaller nanoscale sections produces no significant changes in the anisotropy of Bragg peak profiles. Moreover, under sustained electron irradiation, these domains undergo dramatic rearrangements spanning hundreds of nanometers. Through pinpoint high-dose “impact crater” experiments (19), we also visualize propagating fronts of amorphization extending several hundred nanometers from initial irradiation sites. These results demonstrate that secondary radiolytic processes produce delocalized damage over length scales relevant to macromolecular electron crystallography and single-particle cryogenic electron microscopy (cryoEM) workflows.

## Results

### Protein crystals consist of coherently diffracting domains resembling bend contours

We first conducted a single, low-fluence (0.6 *e*^−^Å^−2^), parallel-beam (0.07 mrad convergence angle) 4D– STEM scan at 85 K on vitrified microcrystals of triclinic lysozyme and monoclinic myoglobin, using a 15 nm full width at half maximum (FWHM) probe. In both cases, we found that each Bragg reflection originates from a small, distinct fraction of the overall coherently diffracting volume (Fig. 2), indicating marked spatial discontinuity among Bragg-active regions. For instance, placing virtual apertures around individual Bragg reflections in lysozyme reveals mostly non-overlapping, roughly parallel CDDs (Fig. 3C-E, H-J). Follow-up experiments on nearby lysozyme crystals replicated this behavior (Fig. S1), with most Bragg reflections originating from flowing swaths representing spatially disconnected subregions of the crystal. Myoglobin, by comparison, exhibits arc-like CDDs thinner and more heterogeneous in shape (Fig. 3M-O, R-T), forming an interlocking pattern. To visualize the contributions from all subregions producing coherent Bragg diffraction at this orientation, we then summed the signal from all selected Bragg peaks to create a composite virtual dark-field (vDF) image (Fig. 3G, Q). We found that the composite vDF image in lysozyme revealed a contiguous diffracting volume (Fig. 3G), whereas in myoglobin the boundaries of individual CDDs remained discernible (Fig. 3Q). Another consistent feature we observed was the spatial proximity between CDDs corresponding to Friedel mates. As shown in Figure S2, five representative pairs of Bragg reflections activated CDDs very close to their symmetry-equivalent counterparts, with several exhibiting partial overlap. In the solid-state TEM literature, similar behavior has been attributed to phenomena called bend contours (25).

**Figure 3.**
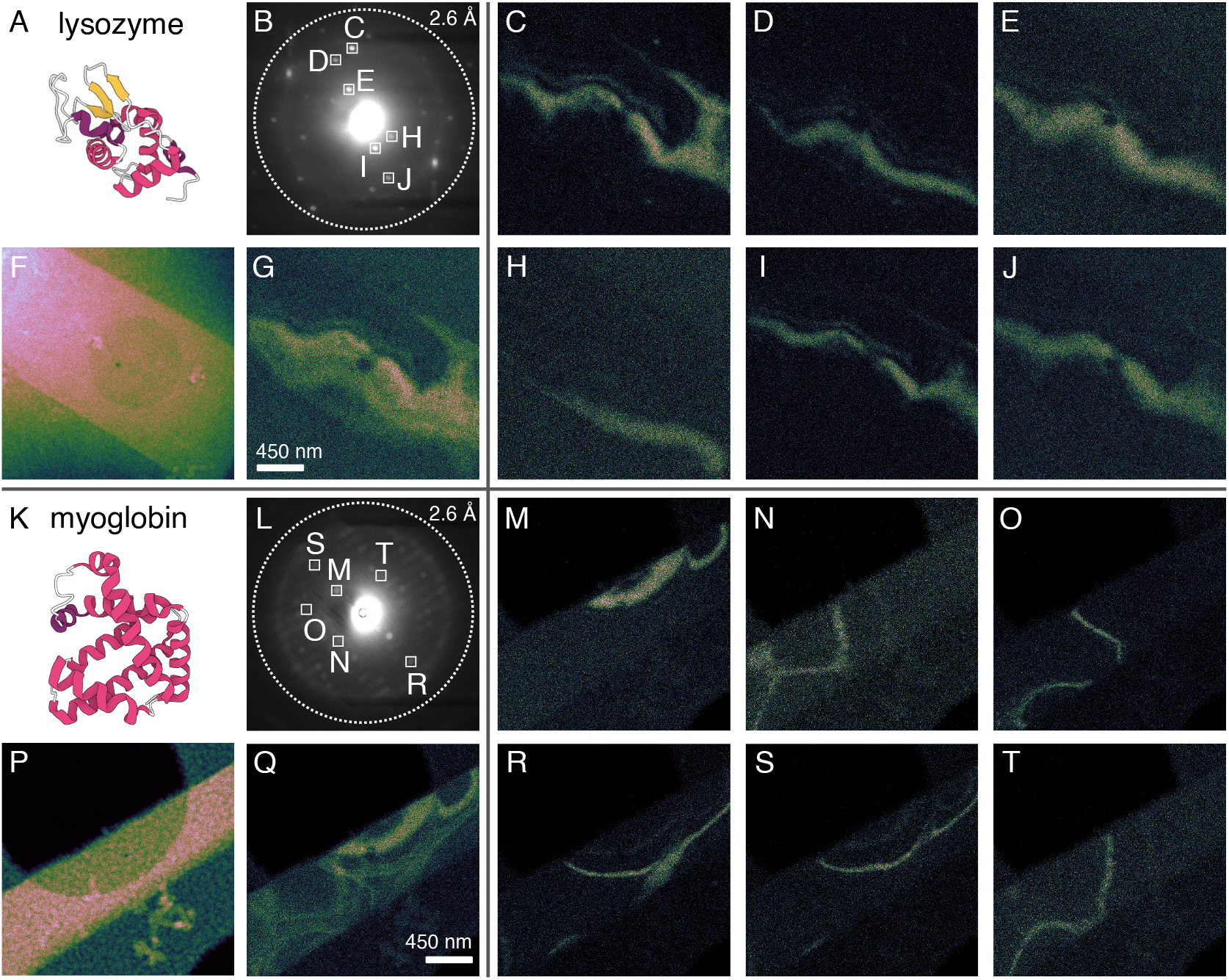
Anatomy of internal protein crystal architecture at the nanoscale. (F, P) High-angle annular dark-field (HAADF-STEM) images of triclinic lysozyme and monoclinic myoglobin microcrystals, respectively, each overlaying a holey carbon support. These images were acquired simultaneously alongside each 4D–STEM scan. (B, L) Selected-area diffraction patterns generated by creating real-space virtual apertures via thresholding the HAADF-STEM images shown in (F, P). White squares correspond to reciprocal-space virtual apertures enclosing specific Bragg reflections. (C-E, H-J) Virtual dark-field images reconstructed from the reciprocal-space virtual apertures in (B). (M-O, R-T) Virtual dark-field images reconstructed from the reciprocal-space virtual apertures in (L). (G, Q) Composite virtual dark-field images summing the Bragg signal in (C-E, H-J) and (M-O, R-T), respectively.

To rationalize the difference in CDD morphology between the two proteins, we note that the triclinic lysozyme polymorph analyzed here lacks any nontrivial crystallographic symmetry operators, whereas the corresponding monoclinic myoglobin polymorph contains a 2_1_ screw axis parallel to the *b* unit cell vector. Protein crystals frequently grow via screw dislocation-mediated spiral mechanisms, as demonstrated by atomic force microscopy (26). We speculate that crystals possessing specific symmetry elements may preferentially nucleate or propagate screw dislocations whose Burgers vectors (27) align with the helical symmetry direction, potentially organizing CDDs into the shapes seen in Fig. 3Q. Furthermore, although several CDDs observed here resemble classical bend contours, the gross morphology of the crystals themselves does not indicate any evidence of forcible mechanical bending (28; 29), suggesting that these internal imperfections could originate during crystal growth.

Next, to probe internal disorder *within* the reconstructed CDDs, we selected a single representative low-resolution reflection in both lysozyme and myoglobin. We then partitioned each CDD into 16 equally sized subregions (Fig. S3A, D), each normalized to have the same total diffracting power. Our goal was to distinguish between two scenarios: a single, continuously monolithic domain versus a collection of smaller, misaligned mosaic blocks whose superposition produces the observed CDD. If the latter were true, subdividing the CDD should yield progressively narrower and sharper Bragg peaks as fewer misoriented blocks contribute to each subregion—i.e., the reciprocal-space signature classically expected from mosaicity (6; 8). To analyze this quantitatively, we fit 2D Gaussian functions to the Bragg reflections arising from each subregion to extract their centroids and standard deviations (*σ*; see SI). We then compared these values to their counterparts obtained from the undivided CDD (Fig. S3B-C, E-F).

Strikingly, the measured *σ* and centroid values remained essentially constant across all 16 subregions, closely matching the value obtained from the full CDD (just under 0.1 mrad; Fig. S3C, F). The individual Bragg peaks also exhibited no significant variations in ellipticity or anisotropy. This invariance suggests that further subdivision does not isolate smaller misoriented constituents—instead, each CDD reconstructed by placing a virtual aperture around a single Bragg reflection already behaves as a monolithic unit. In other words, our procedure effectively isolates what classical theory describes as a single mosaic block. As a caveat, we note that the convergence angles used here (0.07 mrad) limit our sensitivity to subtler rocking-width variations; smaller probes and longer camera lengths would be required to resolve smaller angular spreads (10; 30). Furthermore, we were unable to estimate peak broadening perpendicular to the Ewald sphere surface (e.g., using classical rocking-curve analysis), largely because TEM goniometer imprecision and rapid radiolytic damage preclude such fine-slicing measurements in 4D– STEM.

### Radiolysis causes internal rearrangement of CDDs in protein crystals

As the diffracting power of a crystal fades with accumulating exposure to ionizing radiation, so do the intensities of Bragg reflections. In the aggregate—i.e., the sum of all diffracted intensities—this pattern is generally monotonic. Nevertheless, in some cases, the intensities of individual Bragg peaks can decay nonmonotonically, as observed in synchrotron X-ray studies of protein crystals (31). In previous 4D–STEM work on small-molecule crystals, we showed that vDF movies reconstructed from these nonmonotonically deteriorating reflections reveal CDDs continuously rearranging into morphologies that temporarily better fulfill the Bragg condition *en route* to amorphization (19).

We set out to probe whether crystals of our two model proteins exhibited similar rearrangement behavior. Unlike small-molecule systems, a substantial fraction of the scattering volume of macromolecular crystals consists of disordered solvent (32), typically ranging between 40 and 60%. Furthermore, under vitrified conditions, each crystal is itself embedded in glassy ice. Consequently, we hypothesized that both internal solvent channels and the surrounding vitreous ice matrix could act as radiosensitizers, amplifying the effects of radiolysis by serving as a nearby reservoir of reactive free radicals (particularly H·). This could accelerate radiolytic damage to the point where nonmonotonic decay is too quick to observe. Another salient difference is the looser packing intrinsic to macromolecular self-assembly. To test these hypotheses, we acquired a time series of consecutive 4D–STEM scans on triclinic lysozyme under vitrified conditions at 85 K.

We found that vDF movies reconstructed from nearly any Bragg reflection showed consistent behavior: radiolysis causes CDDs to undergo dramatic internal rearrangement, manifesting as the apparent movement of the CDD across the body of the crystal (Fig. 4). Importantly, this reconfiguration is internal—concurrently acquired high-angle annular dark-field (HAADF–STEM) images (Fig. S4) confirm that there is no accompanying gross translation or rotation of the specimen. In Fig. 4D-E, for instance, the two CDDs travel approximately 500 nm between consecutive scans as their corresponding Bragg reflections decay monotonically. In parallel, the trajectories followed by other CDDs involve marked expansion in their diffracting volumes, leading to nonmonotonic decay patterns (Fig. 4F-G). In these cases, we see fleeting surges in Bragg peak intensity, similar to behavior exhibited by the solvent-free smallmolecule crystals biotin and Ni(dppf)Cl_2_ (19).

**Figure 4.**
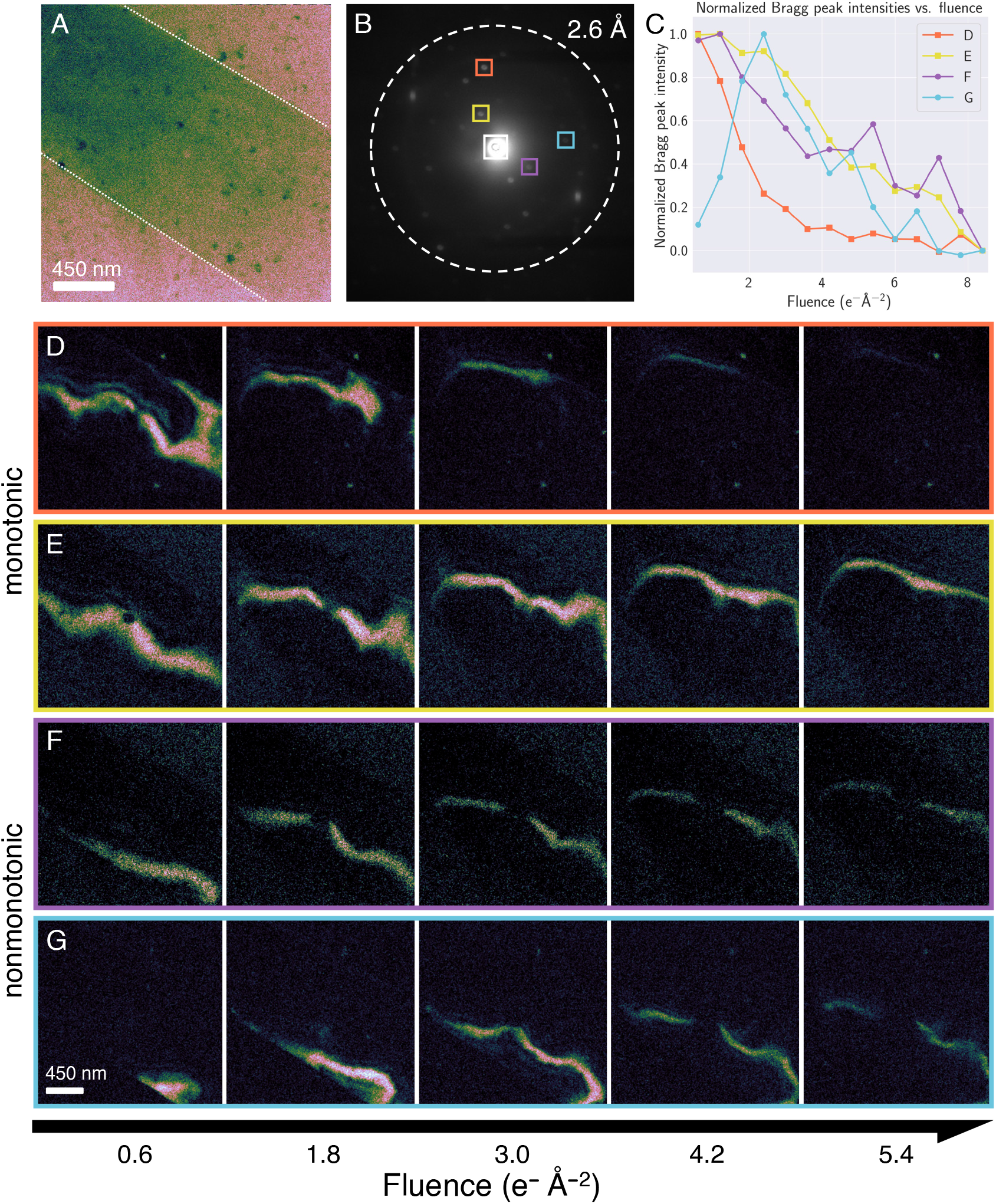
Comparison between virtual dark-field images reconstructed from monotonically and nonmonotonically decaying reflections in vitrified lysozyme at 85 K. (A) Virtual bright-field image of a vitrified lysozyme crystal, reconstructed from the unscattered beam (white box in B). (B) Selected-area diffraction pattern exported from the region outlined in white in (A), showing Bragg reflections extending to 2 Å resolution (dashed circle). Colored boxes indicate virtual reciprocal-space apertures used for tracking intensity decay: monotonic (orange and yellow) and nonmonotonic (purple and blue). (C) Intensities of selected Bragg reflections plotted as a function of accumulating electron fluence, normalized by maximum value. (D, E) Montage of virtual dark-field images reconstructed from the monotonically decaying Bragg reflections marked by the orange and yellow apertures in (B). (F, G) Montage of virtual dark-field images reconstructed from the nonmonotonically decaying Bragg reflections marked by the purple and blue apertures in (B).

The principal distinction, however, lies in the magnitude of the CDD rearrangements observed. Critically, the trajectories of CDD decay in biotin (19) involved comparatively minimal perturbation to initial shape and morphology, with specific CDDs maintaining sharper boundaries and continuously moving during the entirety of the relevant Bragg peak’s lifetime. In comparison, lysozyme CDDs routinely traveled nearly double that distance between consecutive scans, indicating greater fluidity. This discrepancy is striking given that the 4D– STEM experiments in Fig. 4 were acquired at cryogenic temperatures—i.e., conditions under which we might expect some damping of CDD motion—whereas our previous work on biotin was conducted at room temperature. Furthermore, lysozyme and biotin share the same elemental composition, indicating behavior not reflected by traditional metrics such as inelastic cross-sections. In addition to solvent-driven radiosensitization, another possible cause lies in the fundamental mechanical properties of the two systems. Hydrated macromolecular crystals possess elastic moduli around 8× lower than their small-molecule counterparts, largely due to weaker intermolecular bonding and disordered solvent content (33). These intrinsic differences may confer greater flexibility enabling larger-scale CDD rearrangements.

### Tracking distant secondary damage through protein crystals

Previously, we showed that repeated electron irradiation at a single probe position produces expanding fronts of amorphization, which we referred to as “impact craters.” These phenomena represent cascades of secondary radiolytic damage that propagate outward from primary ionization sites (19). We sought to determine whether protein crystals—species far more radiosensitive than the organometallic complexes used in our initial demonstration—exhibit analogous behavior, and whether this sensitivity accelerates crater emergence and expansion. Although macromolecular crystals also decay via radiolysis, the length scales over which these cascades travel in beam-sensitive biological materials remain unexplored. Furthermore, their behavior as a function of temperature is also poorly understood.

To generate the impact crater, we placed the 15 nm probe at the center of a lysozyme crystal and irradiated this dwell point for approximately 200 ms in between consecutive 4D–STEM scans (19) at an incident fluence of 0.4 *e*^−^Å^−2^. Our first foray was conducted at RT. We immediately observed significant delocalization of radiolytic damage, with an impact crater approximately 150 nm in diameter—already 10× bigger than the FWHM of the probe—visible after a single experiment (Fig. 5A, Fig. S5). Diffraction patterns exported from flanking subregions (Fig. 5B) confirmed that Bragg signal was still robust outside the crater (Fig. 5C, E), whereas Bragg reflections within the crater were extinguished (Fig. 5D).

**Figure 5.**
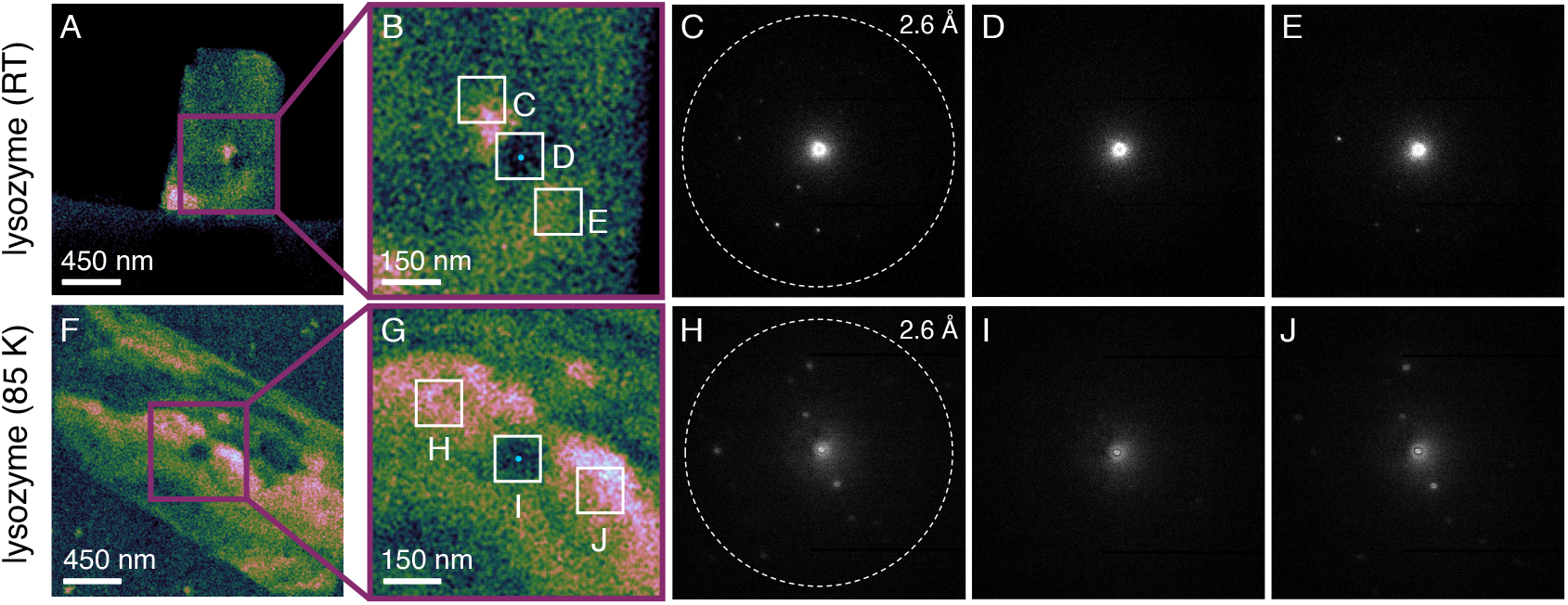
Zoomed-in view of the loss of diffraction signal within the expanding impact crater. (A, F) Composite virtual dark-field images reconstructed from all discernible Bragg reflections. (B, G) Zoomed-in view of the impact crater, with one real-space virtual aperture (D, I) placed directly over the epicenter (cyan dot) and two flanking apertures located approximately 150 nm (C and E) and 200 nm (H and J) away, respectively. The cyan dot represents both the point of incidence and the size of the probe (15 nm at FWHM). Diffraction patterns (C and E, H and J) show the corresponding Bragg signal exported from the labeled regions. In both cases, all Bragg reflections have been extinguished inside the burgeoning crater (D, I), but several remain observable outside its reach (C and E, H and J).

Interestingly, this behavior was replicated under vitrified conditions at 85 K, where a slightly smaller 140 nm crater emerged after a single experiment despite marginally higher fluence (0.6 *e*^−^Å^−2^; Fig. 5F-J). This result was unexpected, since the highly oxidizing hydroxyl radical (·OH)—long considered a major proponent of radiolytic damage at RT (34; 35)—is effectively immobilized below 100 K (36). Therefore, we initially anticipated little to no crater formation under cryogenic conditions due to global suppression of heavier radiolytic fragments such as ·OH. Nevertheless, the comparable level of delocalization at both temperatures implicates a radical species also mobile in ice at 85 K. We propose the hydrogen radical (H·), a well-known byproduct of radiolysis (14; 37). Unlike ·OH, H· faces minimal steric barriers to diffusion, and H_2_ formation via H· radical chemistry is well-documented in vitrified biological specimens under both X-ray and electron irradiation (38; 34; 39; 37). Low-energy secondary electron emission likely also plays some role, although calculations suggest that these species typically thermalize within *<*10 nm boundaries (40; 41) smaller than the probe FWHM. Future experiments using ultralow-loss electron energy-loss spectroscopy in the vibrational regime will test these hypotheses (42).

We then attempted to map the expansion of the nascent impact crater by acquiring consecutive 4D– STEM scans, repeating our pinpoint-irradiation protocol between experiments. Unfortunately, tracking crater growth frame-by-frame was complicated by the scale of the CDD rearrangements described earlier. To image the crater accurately, active CDDs should ideally continue to surround the point of incidence during decay. However, since CDDs in protein crystals travel hundreds of nanometers between consecutive 4D–STEM scans, the underlying mosaic often shifted dramatically before we initiated the next scan, preventing consistent visualization. Furthermore, unlike in Ni(dppf)Cl_2_ (19), virtual dark-field images reconstructed from single Bragg reflections in lysozyme typically did not contain sufficient signal to reconstruct the crater reliably, and several reflections fluctuated in and out of the Bragg condition erratically. This necessitated the usage of composite vDF images to monitor crater growth.

Despite these setbacks, we ultimately collected two dose series—one at RT and one under vitrified conditions at 85 K—in which CDDs serendipitously intersected with the crater site long enough to observe some evolution (Fig. 6). Interestingly, we found that crater expansion did appear to exhibit some temperature dependence, plausibly reflecting the reduced mobility of damaging free radicals at 85 K. At RT, the burgeoning crater doubles in size from approximately 150 nm to 300 nm between the first two frames, whereas the 85 K crater grows more slowly from approximately 140 nm to 200 nm in diameter (Fig. S5). Unfortunately, later scans were difficult to interpret due to the aggressive rate of decay and accompanying CDD rearrangement, particularly at RT. Thus, we hesitate to overinterpret this result. Global diffraction decay was otherwise unremarkable, with Bragg reflections at 85 K lasting approximately 2.5× longer than at RT as expected (Fig. S6).

**Figure 6.**
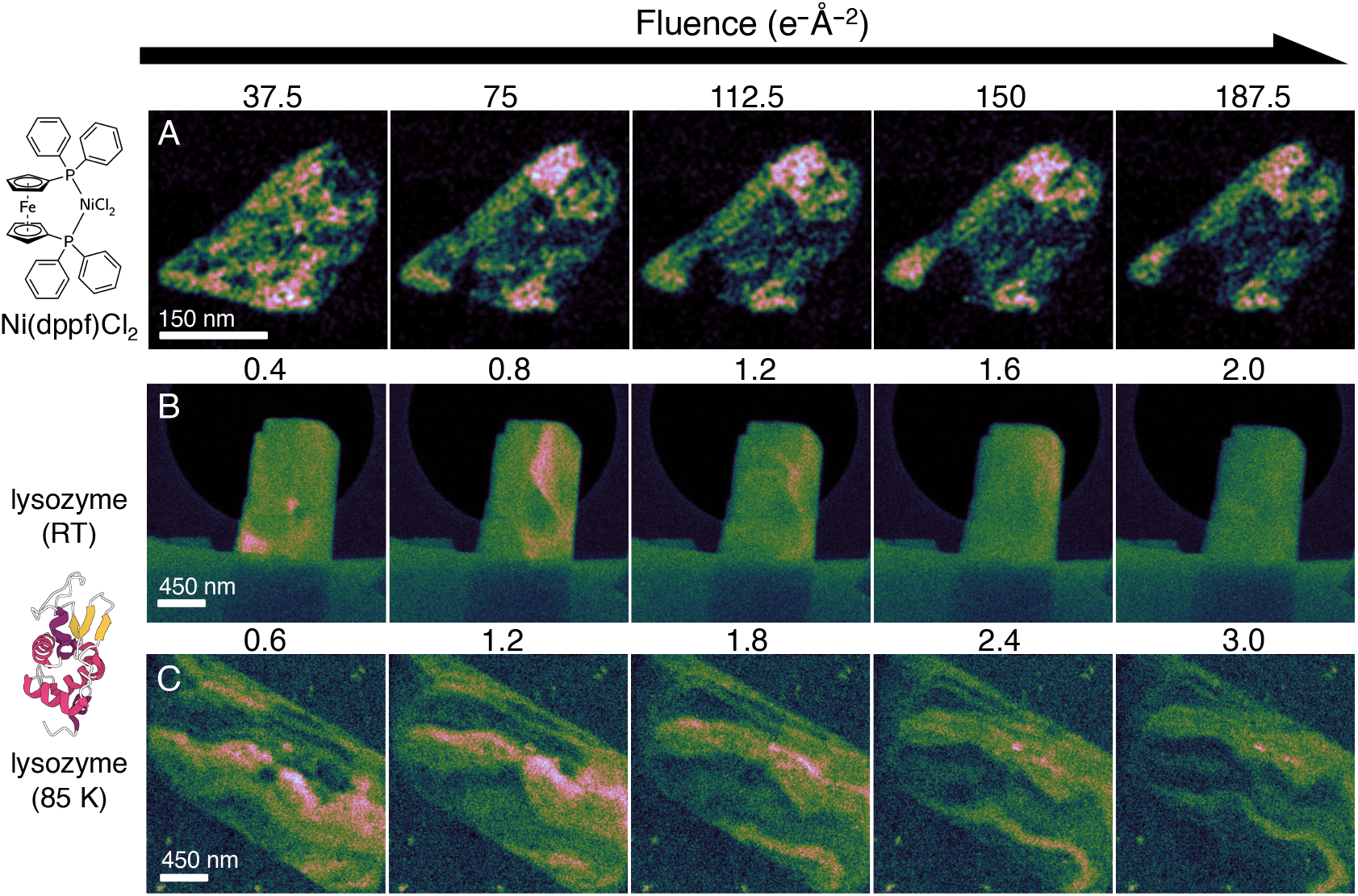
Comparison of impact crater expansion in Ni(dppf)Cl_2_ (19), RT lysozyme, and vitrified lysozyme. (A) Montage of virtual dark-field images reconstructed from a single 4.5 Å reflection in Ni(dppf)Cl_2_ (data taken from (19)), (B) RT lysozyme, and (C) vitrified lysozyme at 85 K. Both lysozyme series consist of composite virtual dark-field images reconstructed from all visible Bragg reflections.

We could, however, draw a much more straightforward comparison with our previous experiments on the organometallic complex Ni(dppf)Cl_2_. Ni(dppf)Cl_2_ required 187 *e*^−^Å^−2^ at 200 kV to produce a 75 nm crater—approximately half the size of the crater seen in Figs. 6B-C and S5. This disparity in delocalization suggests that each primary ionization event in lysozyme produces substantially more secondary radiolytic damage. In radiation chemistry terms, this result is consistent with an elevated *G*-value (43) in lysozyme— i.e., the radiolytic yield of reactive, mobile free radicals per input MGy is likely lower in Ni(dppf)Cl_2_. Interestingly, we also observed that the HAADF-STEM response at the crater site differed strikingly between ambient-temperature and vitrified conditions (Fig. S4). Previously, in Ni(dppf)Cl_2_, repeated irradiation produced a reproducible increase in HAADF intensity at the dwell point, which we attributed to accumulation of carbonaceous contamination (19). Lysozyme at RT exhibited roughly the same behavior, with a slight increase in HAADF signal at the epicenter of the impact crater (Fig. S4A-B). Under vitrified conditions, however, the crater site showed a net loss of HAADF signal, despite comparable cumulative fluence to RT (Fig. S4C-D). Similar behavior was observed in control experiments conducted on pure ice (Fig. S4E).

Two factors could account for these differences. First, Ni(dppf)Cl_2_ is fully aromatic, with every carbon atom belonging to a delocalized *π*-system that absorbs and redistributes excitation energy to confer significant radioresistance (44). Proteins, by contrast, feature predominantly *σ*-bonding: only Phe, Tyr, and Trp possess aromatic sidechains, leaving the majority of the remaining C–H bonds in the polypeptide backbone and other sidechains susceptible to irreversible radiolytic scission and cross-linking. Second, radiolysis of the surrounding ice continuously generates secondary electrons and multiplying free radicals that attack protein molecules, amplifying secondary damage cascades beyond what protein self-ionization would produce. Since HAADF intensity scales with mass-thickness, reduced signal under vitrified conditions suggests local mass loss not present at RT—likely thinning or void formation as highly concentrated radiolysis products escape as volatile species. These observations support our hypothesis that the vitreous ice layer acts as a radiosensitizer.

## Conclusion

Mosaicity has served as an explanatory framework for interpreting the reciprocal-space signatures of crystal imperfection since Darwin (6). Nevertheless, mosaic blocks have remained largely theoretical constructs, with their size, shape, and spatial distribution inferred indirectly from peak profiles and rocking widths. The experiments presented here offer a path toward grounding these models in direct observation. By reconstructing virtual dark-field images from individual Bragg reflections, we visualize protein CDDs whose subdivision produces no further sharpening of peak profiles, suggesting that each domain behaves as an orientationally monolithic unit—effectively a single mosaic block rendered visible. This capability opens the door to mosaicity models parameterized by real-space measurements: domain size distributions and boundary morphologies could all be extracted from 4D–STEM data and used to refine or constrain reciprocal-space fitting. Practically, 4D–STEM could serve as an assay for macromolecular crystal quality, enabling direct comparison of mosaic architecture across crystallization conditions or growth environments.

More broadly, as cryoEM pushes toward higher resolution and faster throughput, characterizing the spatial footprint of radiolytic damage in vitrified solvent will become increasingly important. Our observation that secondary damage propagates hundreds of nanometers from each exposure site suggests that radiolysis may compromise adjacent regions before imaging—an effect unaccounted for in dose management strategies (Fig. S7). Looking forward, chemically informed mitigation strategies that target radical mobility warrant exploration: vitrification in D_2_O, perdeuterated protein expression, or inclusion of radical scavengers could each reduce radiolytic damage propagation while preserving sample integrity. We anticipate that the combination of real-space and reciprocal-space information afforded by 4D–STEM will provide a uniquely powerful lens for dissecting radiolytic damage mechanisms in beam-sensitive biological specimens.

## Methods

### Materials

Hen egg white lysozyme and equine heart myoglobin were purchased from Sigma-Aldrich and used without further purification. Stock solutions of pH 4.5 sodium acetate trihydrate (1.0 M; Hampton Research HR2-789), pH 7.0 sodium acetate trihydrate (4.0 M; Hampton Research HR2-763), sodium nitrate (7.0 M; Hampton Research HR2-661), and ammonium sulfate (3.5 M; Hampton Research HR2-541) were diluted with Milli-Q water to the desired molarity. Sterile 2.0 mm zirconium oxide beads were purchased from Next Advance.

### Crystallization of lysozyme seeds

Crystallization procedures were adapted from literature precedent (45; 46). Hen egg white lysozyme was dissolved in an aqueous solution of 0.05 M sodium acetate trihydrate (pH 4.5) to a concentration of 20 mg mL^−1^ in a

1.0 mL Eppendorf tube. The protein solution was mixed in a 1:1 ratio with a precipitant solution containing 0.4 M sodium nitrate and 0.05 M sodium acetate trihydrate (pH 4.5). The mixture was incubated at 4°C for one day, after which an opaque suspension became visible. The tube was then allowed to rest at room temperature for one week, after which large, colorless crystals were observed at the bottom.

### Crystallization of myoglobin seeds

The crystallization procedure reported by Sherwood *et al*. (47) was followed without modification. Equine heart myoglobin was dissolved in an aqueous solution of 3.0 M ammonium sulfate and 0.1 M sodium acetate (pH 7.0) to a concentration of 8 mg mL^−1^ in a 1.0 mL Eppendorf tube. The tube was left undisturbed at room temperature for one week, after which reddish-brown crystals were observed.

### Crystallization of microcrystals

The procedure described by Hofer *et al*. (48) was adapted. Seed crystals were transferred into a tube containing 5–6 zirconium beads and vortexed in short bursts. In another tube, the solution was mixed at a 2:1:1 ratio with solutions containing modified concentrations of protein and precipitant. For lysozyme, the concentrations of protein and precipitant were doubled to 40 mg mL^−1^ and 0.8 M sodium nitrate, respectively. For myoglobin, the precipitant concentration was tripled while the protein concentration remained unchanged. The mixture was vortexed in bursts, and a cloudy suspension corresponding to micrometer-sized crystals appeared within minutes.

### Grid preparation

The procedure described by Hofer *et al*. (48) was adapted. Quantifoil holey carbon grids were dipped in 1% Tween-20 (Sigma-Aldrich) and then blotted from the metallic side until visibly dry to make the grid hydrophilic. Then, 3 μL of crystal slurry was deposited onto the grid, which was then blotted again from the metallic side until visibly dry. For vitrified samples, the grid was then hand-plunged into liquid ethane and then cryo-transferred into the STEM.

### 4D–STEM data collection

All 4D–STEM experiments were conducted at an accelerating voltage of 300 kV on the double-aberration-corrected Thermo Fisher Titan called TEAM 0.5 located in the National Center for Electron Microscopy facility of the Molecular Foundry at Lawrence Berkeley National Laboratory. Before data acquisition, the incident beam current was reduced to the detection threshold of the fluorescent screen ammeter (<40 pA) by using the monochromator focus as a continuously adjustable gun lens. Beam flux was estimated from previous reference measurements using a Faraday cup and an ammeter under similar conditions. A near-parallel probe was formed by using a custom 10 μm C2 aperture (Norcada) to implement a narrow semiconvergence angle of 0.07 mrad. This produces a beam profile with an approximately 15 nm FWHM. Using a custom DigitalMicrograph script, multiscan 4D–STEM data were acquired with the 4D Camera operating at 87 kHz frame rate. Each scan is composed of 512 × 512 probe positions over a 2.3 μm field-of-view with a probe step size of 4.6 nm.

## Acknowledgments

This work was supported by the following: the National Institutes of Health grants T32GM145388 and R01GM071940; the BioPACIFIC Materials Innovation Platform of the National Science Foundation under award DMR-1933487; the STROBE NSF Science and Technology Center under award DMR-1548924. Work at the Molecular Foundry was supported by the Office of Science, Office of Basic Energy Sciences, of the U.S. Department of Energy under Contract No. DE-AC02-05CH11231. This research used resources of the National Energy Research Scientific Computing Center, a U.S. Department of Energy Office of Science user facility located at Lawrence Berkeley National Laboratory, operated under Contract No. DE-AC02-05CH11231. This work was supported by the Laboratory Directed Research and Development Program of Lawrence Berkeley National Laboratory under U.S. Department of Energy Contract No. DE-AC02-05CH11231. This work was partially funded by the U.S. Department of Energy program “Electron Distillery 2.0: Massive Electron Microscopy Data to Useful Information with AI/ML.” Parts of this manuscript were adapted from the second author’s PhD thesis. The authors thank R. Egerton, D. Cascio, and M. Sawaya for helpful discussions, as well as A. Pattison and T. O’Brien for providing logistical support.

## Data Availability

All 4D–STEM datasets shown here are publicly accessible on Zenodo (49) and viewable using DuSC_explorer (50), our open-source Python-based software.

## Supplementary Information

**Figure S1.**
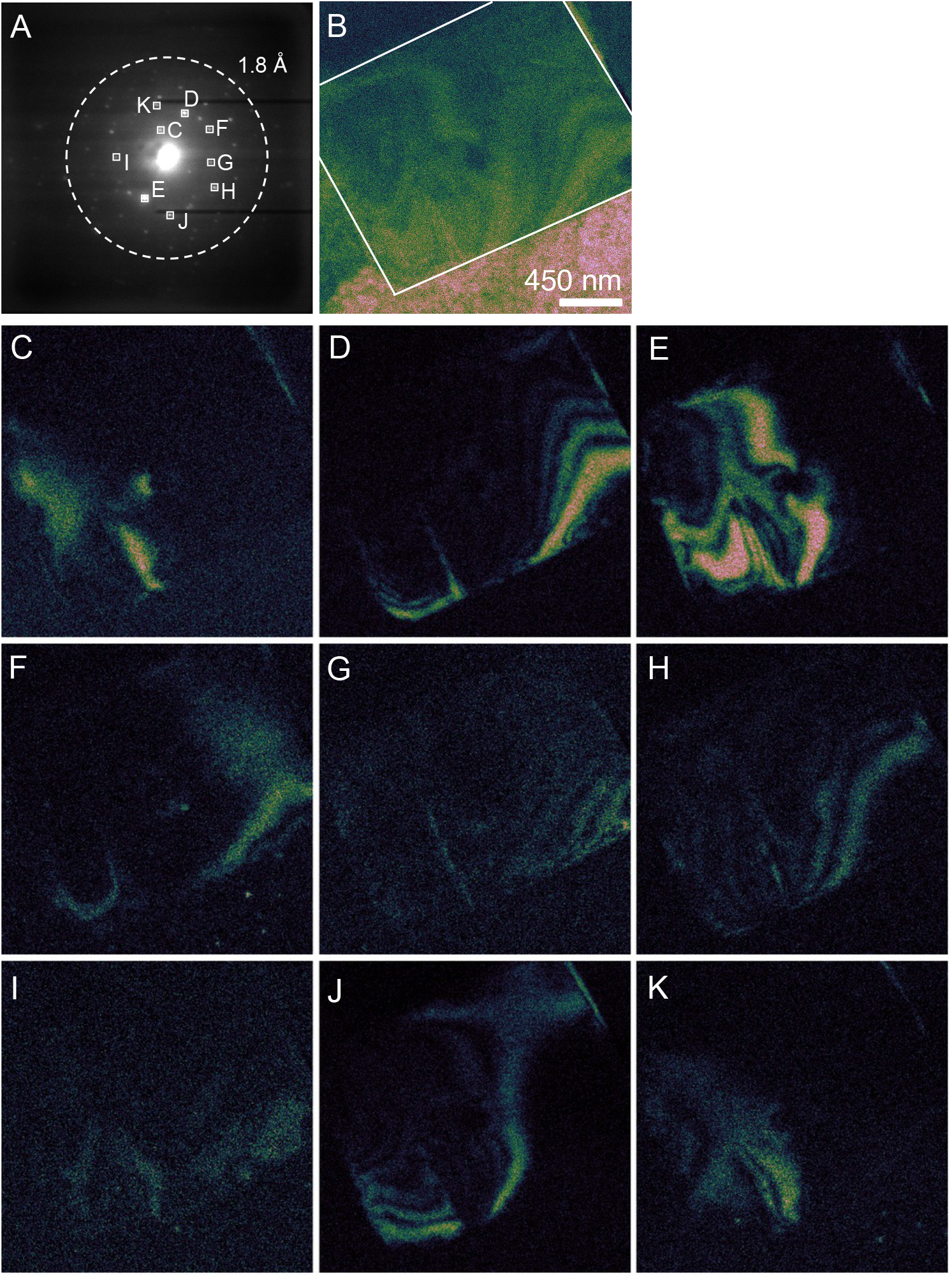
4D–STEM dataset of a lysozyme crystal. The diffraction pattern of the crystal body is shown (A) alongside vDFs of several Bragg reflections. The virtual objective aperture of each reflection is boxed in A and a resolution ring corresponding to a d-spacing of 1.2 Å is shown. The composite image generated by summing vDFs C-K is shown in B, where the crystal body is also outlined by a dashed line.

### Rearrangement of CDDs as a function of fluence

**Figure S2.**
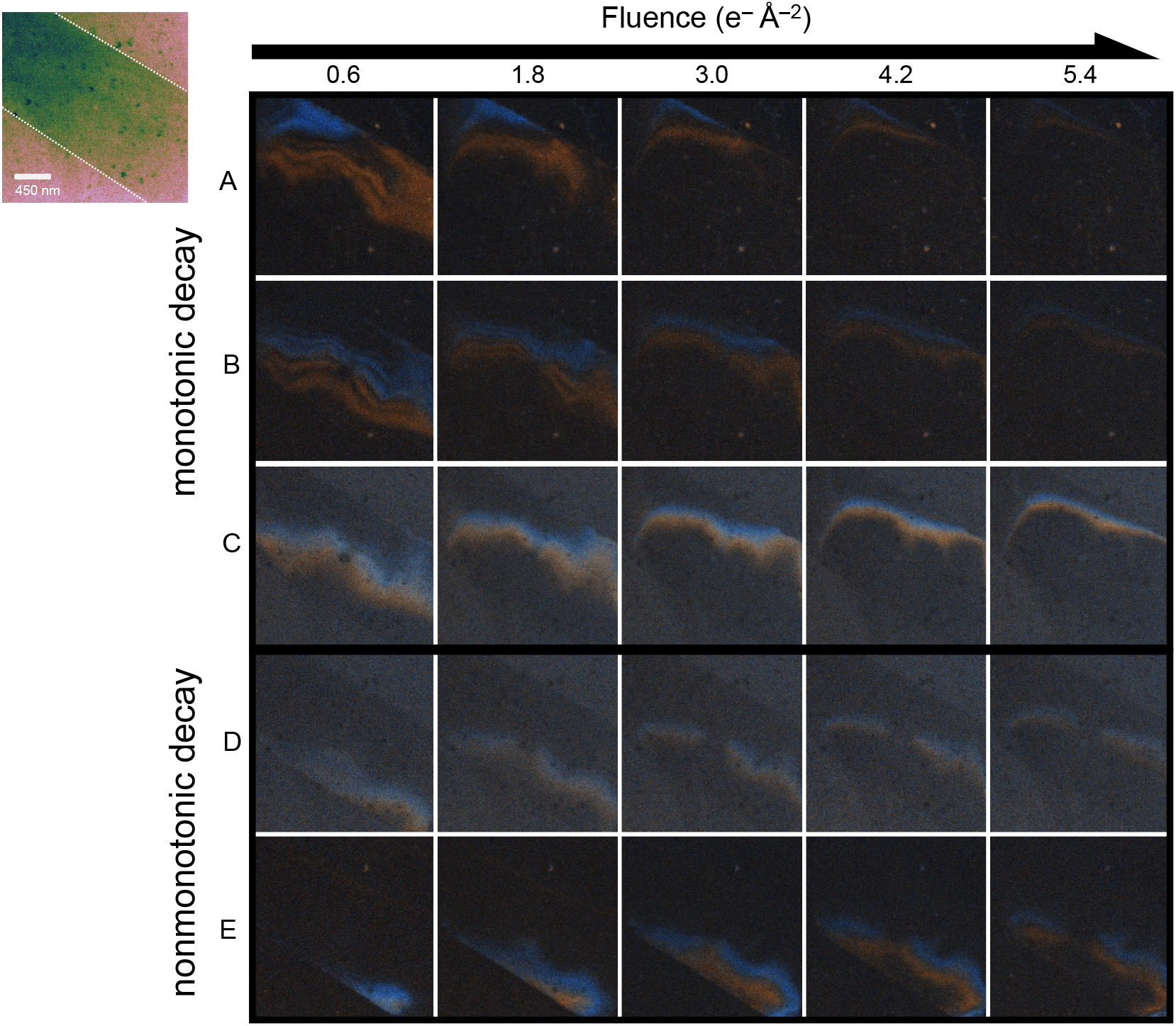
Time series vDFs of lysozyme Friedel pairs. vDF movies of Friedel pairs which exhibit monotonic decay (rows A-C) and nonmonotonic decay (rows D and E). The CDDs of each Bragg peak of the Friedel pair are highlighted in red and blue. A vBF image of the crystal is shown at the top-left, with a dashed-white outline of the body of the crystal.

**Figure S3.**
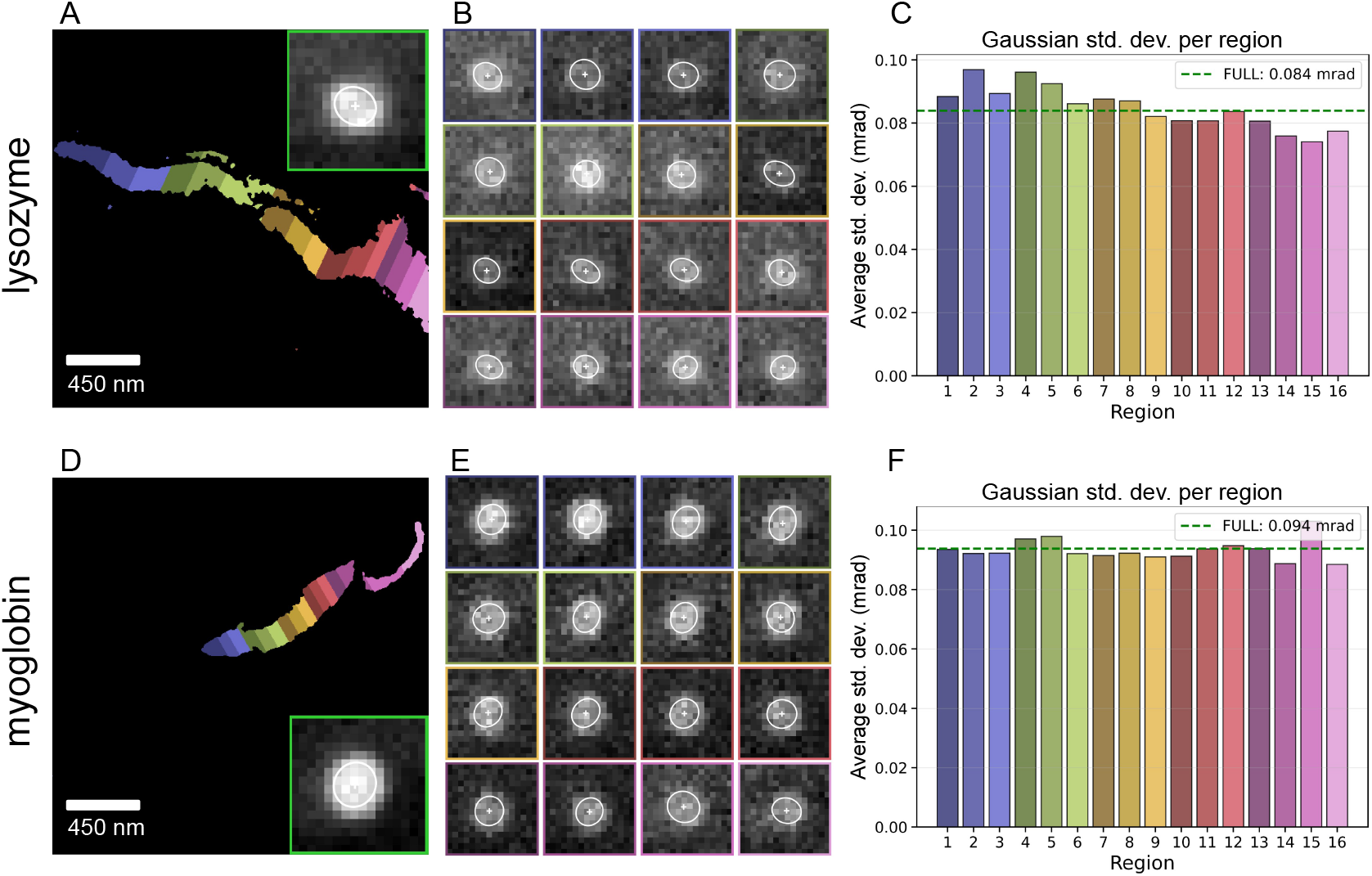
Anatomy of a coherently diffracting domain. (A, D) The CDDs reconstructed in Fig. 3I and J are subdivided into 16 subregions of equal diffracting power. Inset: the corresponding Bragg reflection produced when signal is exported from all 16 subregions combined. (B, E) Coherent Bragg signal corresponding to each individual subregion depicted in (A, D). Each reflection is annotated with an ellipsoid demarcating one standard deviation (*σ*) radius from the centroid. (C, F) Comparison of the average *σ* for each peak in (B, E), suggesting no significant broadening indicative of classical mosaicity. All peaks were cropped using the same pixel values from the original diffraction pattern; in other words, they were not cropped around each of their respective peak centroids. The mosaic block size of the myoglobin reflection and CDD is estimated, using Scherrer’s equation (1), to be larger than 72 nm.

### Gaussian analysis of sub-CDDs

To analyze the Bragg peak characteristics, the reflections were cropped from the scan. A vDF of the reflection was generated and manually low-pass filtered and thresholded to produce a mask which only included the CDD, eliminating noise or other sources of signal (e.g., ice crystals). The masked vDFs were then split into 16 sub-CDDs of equal electron counts (within ∼2%). Each of these sub-CDDs was then turned into its own mask for producing the diffraction pattern from each of those subregions. These resultant Bragg reflections were modeled with a 2D Gaussian function for further analysis.

Areas and eccentricities were determined for the ellipse formed from one continuous standard deviation (*σ*) around the Gaussian function’s centroid. The area was calculated as:

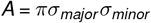

The eccentricity was then calculated as:

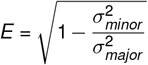

Where *σ*_*major*_ and *σ*_*minor*_ are the standard deviations along the major and minor axes. Values of *E* ≈ 0 correspond to near-perfect circles, and peaks with eccentricity closer to 1 are more ellipsoidal.

### Estimation of mosaic block size

The minimum mosaic block size within the crystal was estimated using the Scherrer equation (1), which relates the angular broadening of a diffraction peak to its corresponding crystallite size. The Scherrer equation is given by

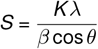

where *S* is the crystallite size, *K* is a shape factor (taken here as 0.95), *λ* is the electron wavelength, *β* is the full width at half maximum (FWHM) of the diffraction peak in radians, and *θ* is the Bragg angle. For the small scattering angles typical of electron diffraction, cos *θ* ≈ 1.

The angular scale at the detector is determined by the experimental geometry. For small angles, the Bragg angle is related to the radial detector position by *θ* = *r/*2*L*, where *r* is the distance from the direct beam and *L* is the detector distance. With a pixel size of 0.01 mm and *L* = 135 mm, this corresponds to 0.037 mrad per pixel.

Analysis of the diffraction peak widths revealed that the Bragg reflections and direct beam exhibited nearly identical angular widths. Assuming the diameters are the same to within 10% (0.7 pixels), this corresponds to an upper limit on the size-related broadening of

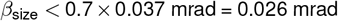

Substituting this upper bound into the Scherrer equation yields a lower limit on the mosaic block size:

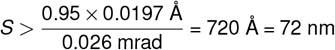

This result indicates that the mosaic blocks are *at minimum* 72 nm in size, comparable to the overall crystal dimensions observed in electron microscopy. The absence of detectable peak broadening suggests that the crystal is composed of relatively large, coherently diffracting domains, with any mosaicity below the resolution limit of the present experimental configuration.

**Figure S4.**
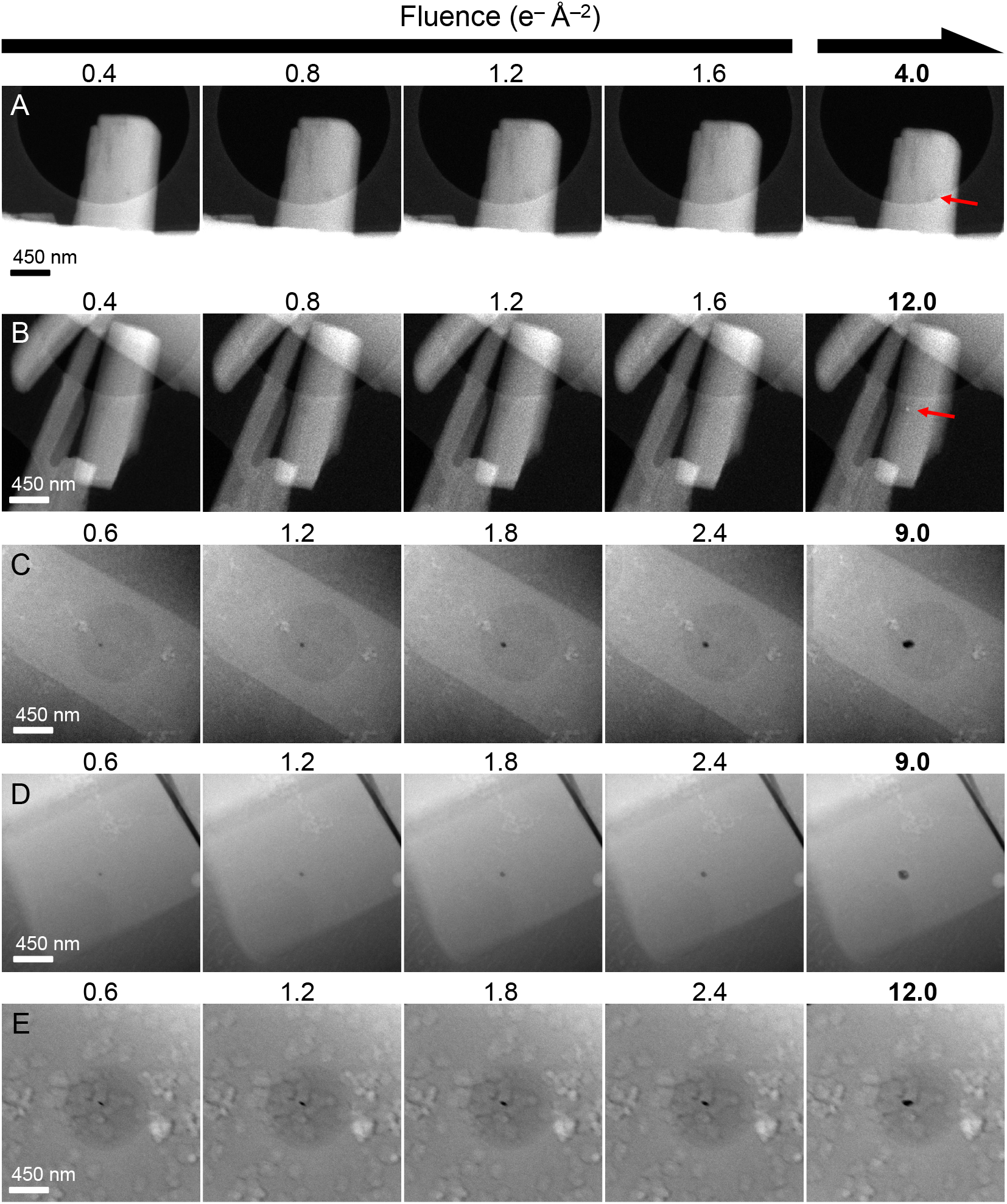
HAADF-STEM images from impact crater series. HAADF-STEM image series of lysozyme at room temperature (A, B) and ∼85 K (C, D) as well as a series of ice at ∼85 K (E). The ice sample is comprised of a layer of vitreous ice along with some visible ice crystal domains. In between each image, there is an extra dose applied at or near the center of the frame, leading to changes in HAADF contrast. The first four images in each series are the first four images taken, whereas the last column of images displays the last image taken of the series, regardless of how many further images were collected; this is also shown by the fluence labels above each image. In the final frame of the room temperature experiments there is a red arrow indicating the location of the impact crater epicenter; in the 85 K samples, it is visible as the growing dark circle.

**Figure S5.**
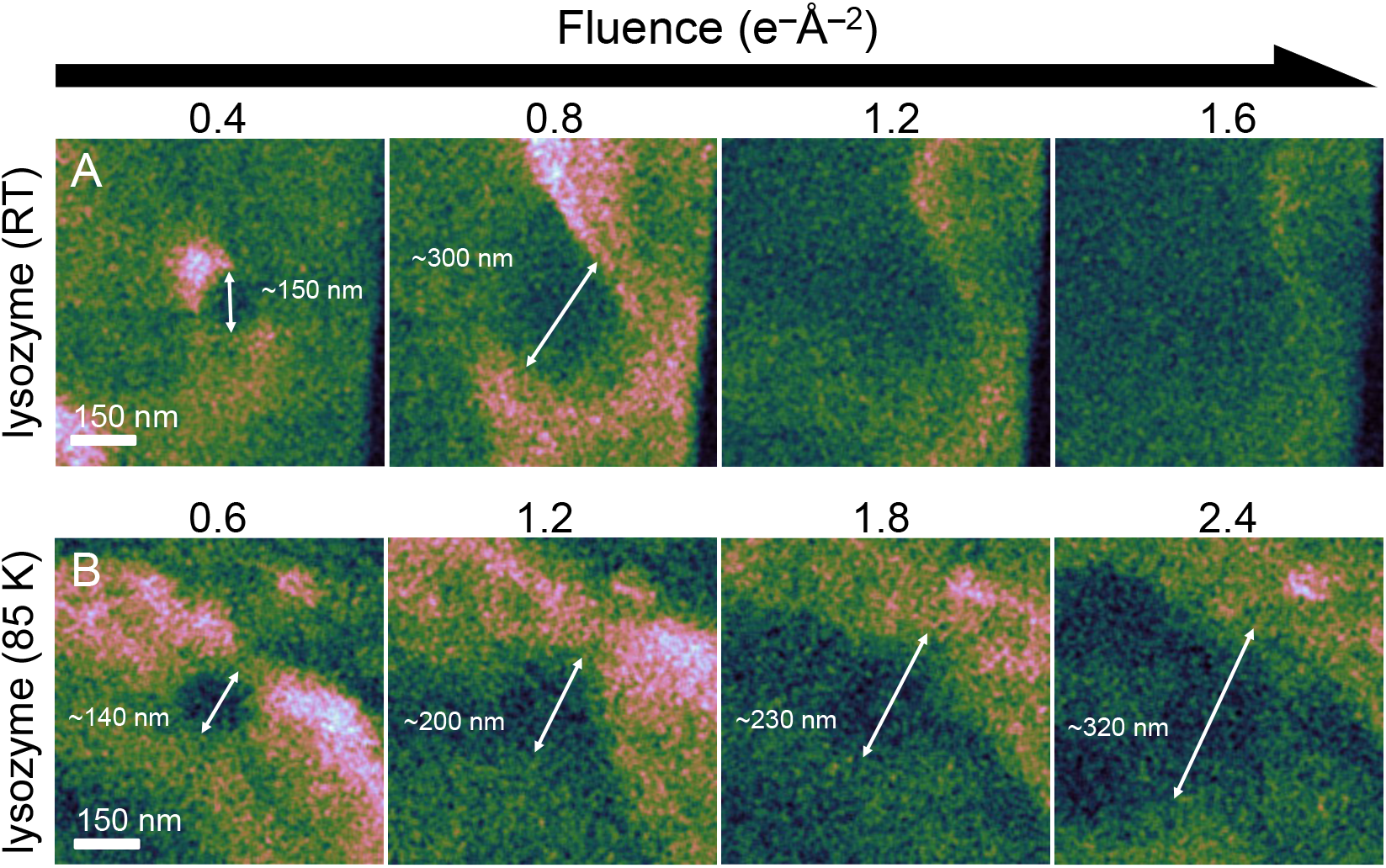
Tracking the growth of the impact crater in lysozyme at room temperature and 85 K. Zoomed-in vDF images of samples shown in Figure 6B and C. An estimated measurement of the crater diameter is drawn where visible.

**Figure S6.**
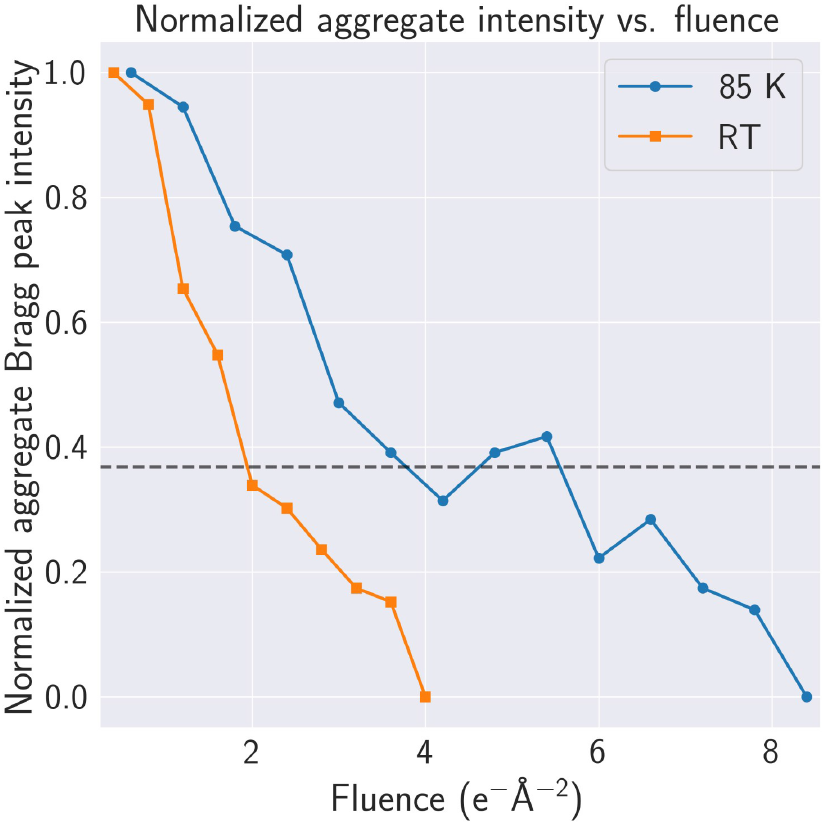
Temperature dependence of diffraction lifetime during 4D–STEM scanning. Normalized aggregate Bragg peak intensity (aggregate of measured Bragg-peak intensities per scan, normalized to the first scan) plotted as a function of accumulated incident fluence for lysozyme microcrystals imaged at 85 K (blue) and room temperature (orange). The horizontal dashed line marks the *I/I*_0_ = 1*/e* criterion used to estimate the characteristic fluence-to-decay, highlighting the slower intensity loss at cryogenic temperature under otherwise comparable acquisition conditions.

**Figure S7.**
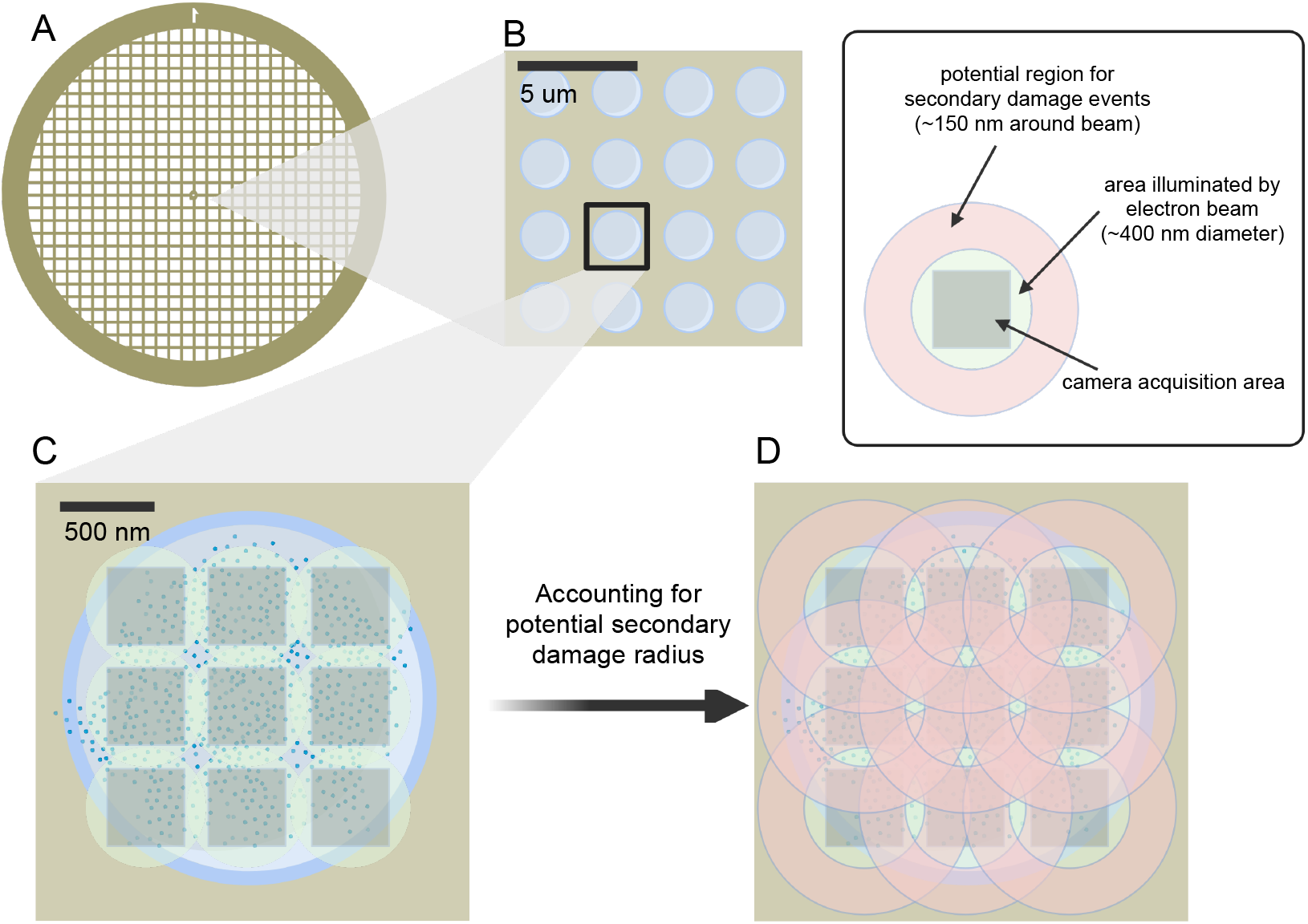
Potential effects of distant secondary damage to cryoEM imaging workflows. Holey TEM grids (A) used in cryoEM have an array of holes (B). Within each hole, an array of imaging targets is selected (C, imaged area is shaded black, area illuminated by the beam is shaded green). However, secondary radicals may be propagating from one illuminated area to a future target (D, shaded red). The Krios G4 (Thermo Fisher Scientific) with fringe-free imaging can form an electron beam of ∼400 nm diameter. This enables higher throughput imaging as imaging targets may be placed closer together without irradiating regions to be imaged next. However, the impact crater in lysozyme is roughly 150 nm after one scan, subsequently growing even larger. This suggests that a cascading form of radiation damage may lead to some amount of “pre-dose” with imaging targets placed so closely together. *Created in BioRender. Nia, S. (2026) https://BioRender.com/0g2pes9*

## Notes

https://doi.org/10.5281/zenodo.18200830

### Competing Interest Statement

The authors have declared no competing interest.

### Summary of Updates

Minor typos corrected in the text. Minor changes to graphics in figures.

